# Integrating identification and quantification uncertainty for differential protein abundance analysis with Triqler

**DOI:** 10.1101/2020.09.24.311605

**Authors:** Matthew The, Lukas Käll

## Abstract

Protein quantification for shotgun proteomics is a complicated process where errors can be introduced in each of the steps. Triqler is a Python package that estimates and integrates errors of the different parts of the label-free protein quantification pipeline into a single Bayesian model. Specifically, it weighs the quantitative values by the confidence we have in the correctness of the corresponding PSM. Furthermore, it treats missing values in a way that reflects their uncertainty relative to observed values. Finally, it combines these error estimates in a single differential abundance FDR that not only reflects the errors and uncertainties in quantification but also in identification. In this tutorial, we show how to (1) generate input data for Triqler from quantification packages such as MaxQuant and Quandenser, (2) run Triqler and what the different options are, (3) interpret the results, (4) investigate the posterior distributions of a protein of interest in detail and (5) verify that the hyperparameter estimations are sensible.

## 1 Introduction

Shotgun proteomics has proven to be a useful technique for identifying and quantifying proteins across multiple samples. The identification process of fragment mass spectra has received ample scrutiny in terms of statistical rigor, with target-decoy analysis [1] and false discovery rates (FDRs) [2] being widely adopted by the field. In contrast, as a result of its complexity, error estimation in the quantification process is often limited to a sequence of, frequently heuristic, thresholds. Application of these thresholds is often well-intended and does indeed eliminate false discoveries, but often fails to take into account biases and interactions these create with other thresholds. As a prime example, consider the common practice of applying a fold change threshold after obtaining a list of differentially abundant proteins by a *t*-test. Whereas the original list of differentially abundant proteins was controlled at, say, 5% FDR, there are no guarantees that this FDR is valid for the list after application of the fold change threshold, as there could well be an enrichment of false positives at higher fold changes.

To make matters even more complicated, missing value imputation is an essential part of data analysis for label-free quantification, as often more than 50% of the data is missing due to the stochastic nature of data-dependent acquisition. These imputation methods can introduce many different types of errors [3, 4] which in turn can have a big impact on the statistical test employed for differential abundance testing.

To address these issues, we developed Triqler [5], a Python package that uses a Bayesian model to integrate errors at the different levels of protein quantification. Specifically, it combines identification and quantification errors into a single differential abundance FDR. The Bayesian model propagates and integrates the uncertainties from feature, PSM, peptide, protein and treatment group level to a final posterior distribution of the fold change between two treatment groups. This is especially useful for missing values, for which we can specify a probability distribution over a range of likely values with a higher likelihood towards lower values. In this way, missing values are considered less reliable than observed values, as is intuitively clear.

The input format for Triqler is a simple tab-delimited file (*see* Subheading 2.6) and can most easily be obtained by converting output files from MaxQuant [6] (*see* Subheading 3.1) or Quandenser [7] (*see* Subheading 3.2). Alternatively, one can use any search engine of choice and add quantification information using Dinosaur [8] (*see* Subheading 3.3). For this tutorial, we explain how to run Triqler (*see* Subheading 3.4), what the individual steps inside Triqler are (*see* Subheading 3.5) and how to interpret the output (*see* Subheading 3.6). Finally, we dive deeper into the results by looking at posterior distributions for a protein of interest (*see* Subheading 3.7) and the hyperparameter estimation (*see* Subheading 3.8).

## 2 Material

### 2.1 Requirements

Triqler can be run on any system with Python 2 or 3 installed and has been tested on Windows, Mac OS X and Linux. For typical datasets, only a modest amount of RAM is required (<2GB) and can, thus, be run on any desktop computer. To accommodate large-scale dataset analysis, Triqler also supports multicore processing.

For users unfamiliar with the Python environment, we recommend installation of Python through the Anaconda environment, freely available for all major operating systems from https://www.anaconda.com/products/individual.

### 2.2 Software install

Triqler is available through the pip command:

~~~
   $ pip install triqler
~~~

Alternatively, you can clone the GitHub repository and build the package locally:

~~~
   $ git clone https://github.com/statisticalbiotechnology/triqler.git
   $ cd triqler
   $ pip install.
~~~

These commands will download the latest release of Triqler (currently v0.5). To update the package, replace `$ pip install by $ pip install --upgrade` in either command.

### 2.3 Datatype

Triqler requires a list of PSMs, as well as corresponding information regarding intensity, sample and experimental condition. Unfortunately, not all search engines provide intensity information with their PSMs and this information might, thus, have to be added in a separate step (*see* Subheading 3.3).

### 2.4 Data size — Number of samples

As Triqler aims to determine differential abundance, it requires at least 2 experimental conditions (e.g. case and control) and 3 biological or technical replicates per condition.

### 2.5 Data size — Number of proteins

The parameter estimation of the error model of Triqler depends on a background distribution of proteins that are not differentially abundant. Therefore, we recommend that the software should only be used in situations where the majority of the proteins (> 70%) are expected not to be differentially abundant between the conditions and more than 100 proteins are present (*see* **Note 1**).

### 2.6 Input format

To simplify the creation of input files, we provide a set of converters that take results from packages such as MaxQuant (*see* Subheading 3.1), Quandenser (*see* Subheading 3.2), Dinosaur (*see* Subheading 3.3) and Tide (*see* Subheading 3.3) search and convert them to the Triqler input format.

Alternatively, it is not too complicated to generate a Triqler input file by yourself. An example input file is provided in the GitHub repository at https://github.com/statisticalbiotechnology/triqler/blob/master/example/iPRG2016.tsv. We will use this example file throughout this chapter to demonstrate several characteristics of Triqler.

The input format consists of 7 columns separated by tabs (see Fig. 1), indicated by the following headers: `run, condition, charge, searchScore, intensity, peptide, proteins`. Each row consists of a PSM result from a search engine of choice (`charge, searchScore, peptide, proteins`), together with the intensity from the corresponding MS1 feature (`intensity`) and meta-information regarding the sample the PSM corresponds to (`run, condition`). If a peptide is shared between multiple proteins, the protein names are separated by semicolons (*see* **Note 2**).

**Fig. 1.**
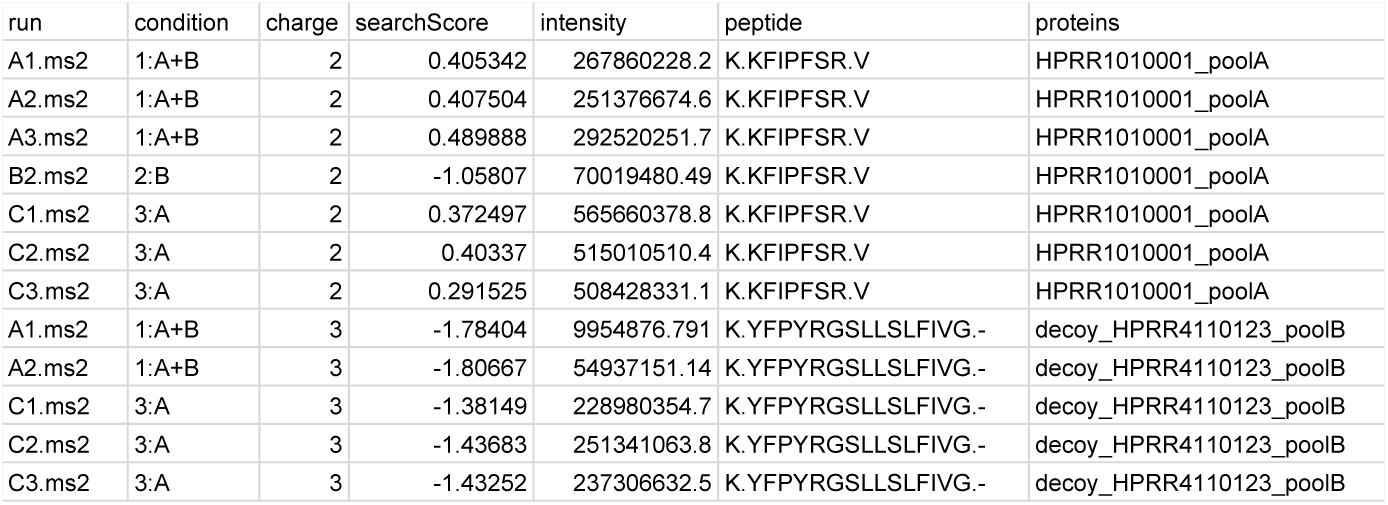
An example of the Triqler input format.

## 3 Methods

As the input to Triqler is unfortunately rather non-standard, we start with 3 different methods of generating an input file from existing pipelines. This will be followed by running Triqler itself and interpreting the results, as well as diving deeper into the results for particular proteins of interest.

Commands on the command line are prepended by the $ character (e.g. `$ python --version`), which should be omitted when actually running the command (e.g. `python --version`). Command line inputs are written between the < and > symbols, e.g. `<X>`, which should be replaced by the user by the relevant value or file path.

### 3.1 Generating Triqler input from MaxQuant

Triqler contains functionality to convert a MaxQuant evidence.txt output file into a Triqler input file [9] (an example is given below, *see* Subheading 3.1, **Step 3**):

~~~
     $ python -m triqler.convert.maxquant --file_list_file <L>
     --out_file <OUT_FILE> <IN_FILE>
~~~

Explanation of the command line parameters:

~~~
     <IN_FILE>
~~~

the file called `evidence.txt` present in the `combined/txt` folder of the MaxQuant results.

~~~
     --file_list_file <L>
~~~

a simple tab-separated text file with spectrum file names in the first column and treatment group in the second column (see Fig. 2, left). The spectrum file names should be the same as in the `Raw file` column name in `evidence.txt` without the preceding path. In the case of fractionated samples, the third and fourth columns should contain the sample name and fraction respectively (see Fig. 2, right).

~~~
     --out_file <OUT>
~~~

the output of this converter, a Triqler input file.

**Fig. 2.**
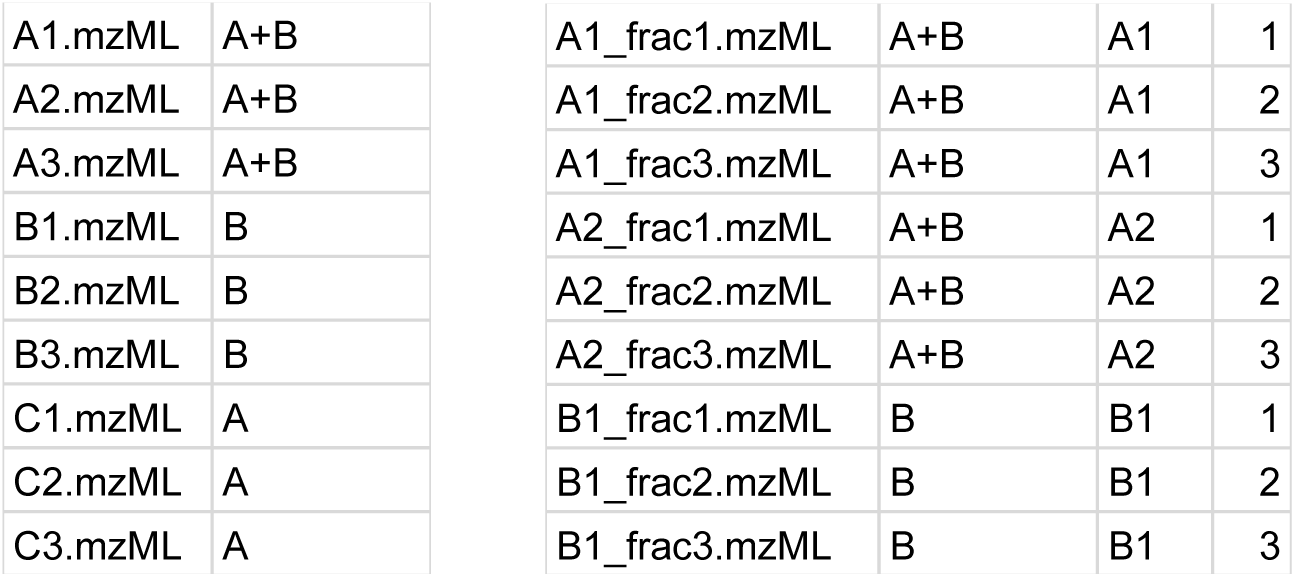
An example of the Triqler file list input format. The simple format (left) only contains the file name and experimental condition. The extended format (right) also includes columns for the sample name and fraction number.

Below are the step-by-step instructions to process a MaxQuant `evidence.txt` file with Triqler:

1. For best performance, we recommended running MaxQuant by setting all FDR thresholds (PSM, peptide and protein-level) to 100% (*see* **Note 3**) and turning off matches-between-runs (*see* **Note 4**).
2. Create a tab-delimited file with the file metadata as described above, e.g. `file_metainfo.txt`.
3. Run the MaxQuant converter:

~~~
  $ python -m triqler.convert.maxquant --file_list_file
  file_metainfo.txt --out_file triqler_input.tsv evidence.txt
~~~
4. MaxQuant uses the decoy prefix REV__ by default, so remember to change this prefix accordingly:

~~~
  $ python -m triqler --decoy_pattern REV  triqler_input.tsv
~~~

### 3.2 Generating Triqler input from Quandenser

Quandenser is a peptide quantification package that employs unsupervised clustering on both MS1 and MS2 data, without assigning peptide sequences. The benefit is that one can run the identification part as often as desired, without having to redo the quantification. The current Quandenser converter relies on that the search results are post-processed by Percolator [10, 11].

The interface for this converter is as follows (an example is given below, *see* Subheading 3.2, **Step 4**):

~~~
    $ python -m triqler.convert.quandenser --file_list_file <L>
    --psm_files <TARGET>,<DECOY> --out_file <OUT> <IN_FILE>
~~~

Explanation of the command line parameters:

~~~
    <IN_FILE>
~~~

the file called `Quandenser.feature_groups.tsv` in the Quandenser results folder.

~~~
    --file_list_file <L>
~~~

a simple tab-separated text file with spectrum file names in the first column and condition in the second column. In the case of fractionated samples, the third and fourth columns should contain the sample name and fraction respectively (see Fig. 2).

~~~
    --psm_files <TARGET>,<DECOY>
~~~

the target/decoy PSM output files from Percolator, separated by commas. Both output files from stand-alone Percolator as well as from `crux percolator` are supported (*see* **Note 5**).

~~~
    --out_file <OUT>
~~~

the output of this converter, a Triqler input file.

Below are the step-by-step instructions to process the Quandenser output files with Triqler (*see* **Note 6**).

1. Run Quandenser on your input files. This will produce a file called `Quandenser.feature_groups.tsv` as well as one or more consensus spectra files in the `consensus_spectra` folder.
2. Search all consensus spectra files with a search engine of choice, preferably one supported by Percolator (*see* **Note 7**).
3. Run Percolator on the search results files, make sure to use both the `--results-psms` and `--decoy-results-psms` to obtain results on PSM level with both targets and decoys reported.
4. Convert the search results

~~~
  $ python -m triqler.convert.quandenser --file_list_file
  file_metainfo.txt --out_file triqler_input.tsv
  --psm_files percolator.target_psms.txt.percolator.decoy_psms.txt
  Quandenser.feature_groups.tsv
~~~
5. Run Triqler:

~~~
  $ python -m triqler triqler_input.tsv
~~~

### 3.3 Generating Triqler input from Dinosaur

Triqler also provides functionality to do quantitative analysis on search results from a search engine of choice, post-processed by Percolator (*see* **Note 8**), by using Dinosaur to add MS1 quantification information to the MS2 spectra.

The interface for this converter is as follows (an example is given below, *see* Subheading 3.3, **Step 4**):

~~~
    $ python -m triqler.convert.dinosaur --file_list_file <L>
    --psm_files <TARGET>,<DECOY> --out_file <OUT> <IN_FILES>
~~~

The command line parameters are the same as in Subheading 3.2, with the exception that the input files now are feature-to-spectrum mapping files, most easily produced by using our Dinosaur adapter for Python (*see* **Note 9**).

Below are the step-by-step instructions to obtain quantification information with Triqler using a search engine of choice. (*see* **Note 10**).

1. Run the Dinosaur adapter for Python on your input mzML files (*see* **Note 11**). This will produce several files named `dinosaur/<file_name>.feature_map.tsv`, and, optionally, spectrum files named `dinosaur/<file_name>.recalibrated.<spectrum_format>`.
2. Search your input mzML files or the recalibrated MS2 spectrum files (*see* **Note 12**) with a search engine of choice, preferably one supported by Percolator (*see* **Note 13**).
3. Run Percolator on the search results files, make sure to use both the `--results-psms` and `--decoy-results-psms` to obtain results on PSM level with both targets and decoys reported.
4. Convert the search results

~~~
  $ python -m triqler.convert.dinosaur --file_list_file
  file_metainfo.txt --out_file triqler_input.tsv
  --psm_files percolator.target_psms.txt.percolator.decoy_psms.txt
  dinosaur/*.feature_map.tsv
~~~
5. Run Triqler:

~~~
  $ python -m triqler triqler_input.tsv
~~~

### 3.4 Triqler interface

To verify that Triqler was installed correctly and to get an overview of the command line parameters, run the following command:

~~~
    $ python -m triqler --help
~~~

This should produce the following help text. The individual command line arguments are explained further below:

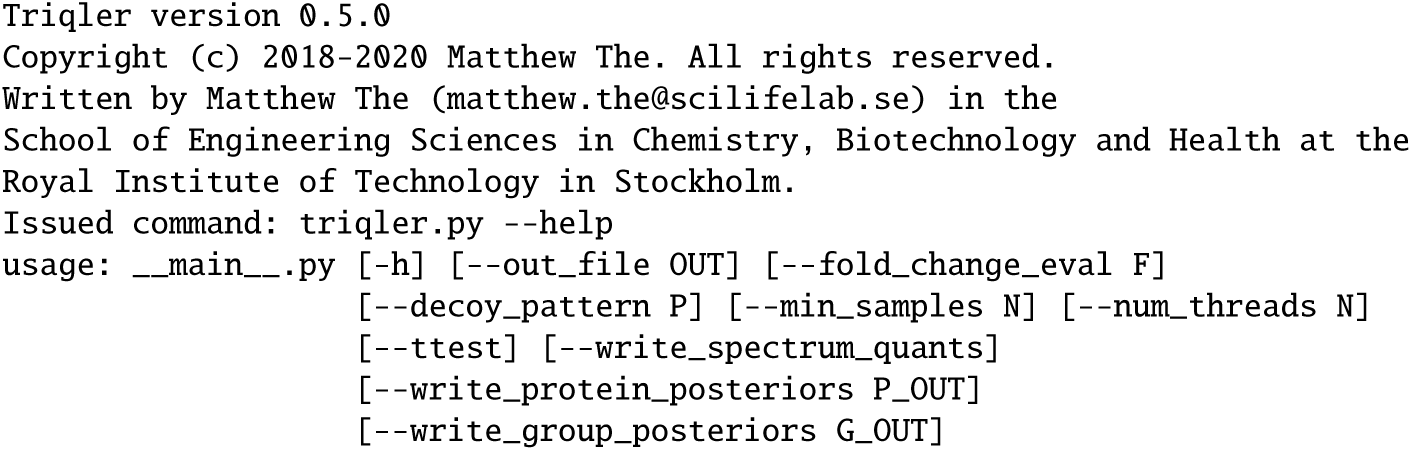
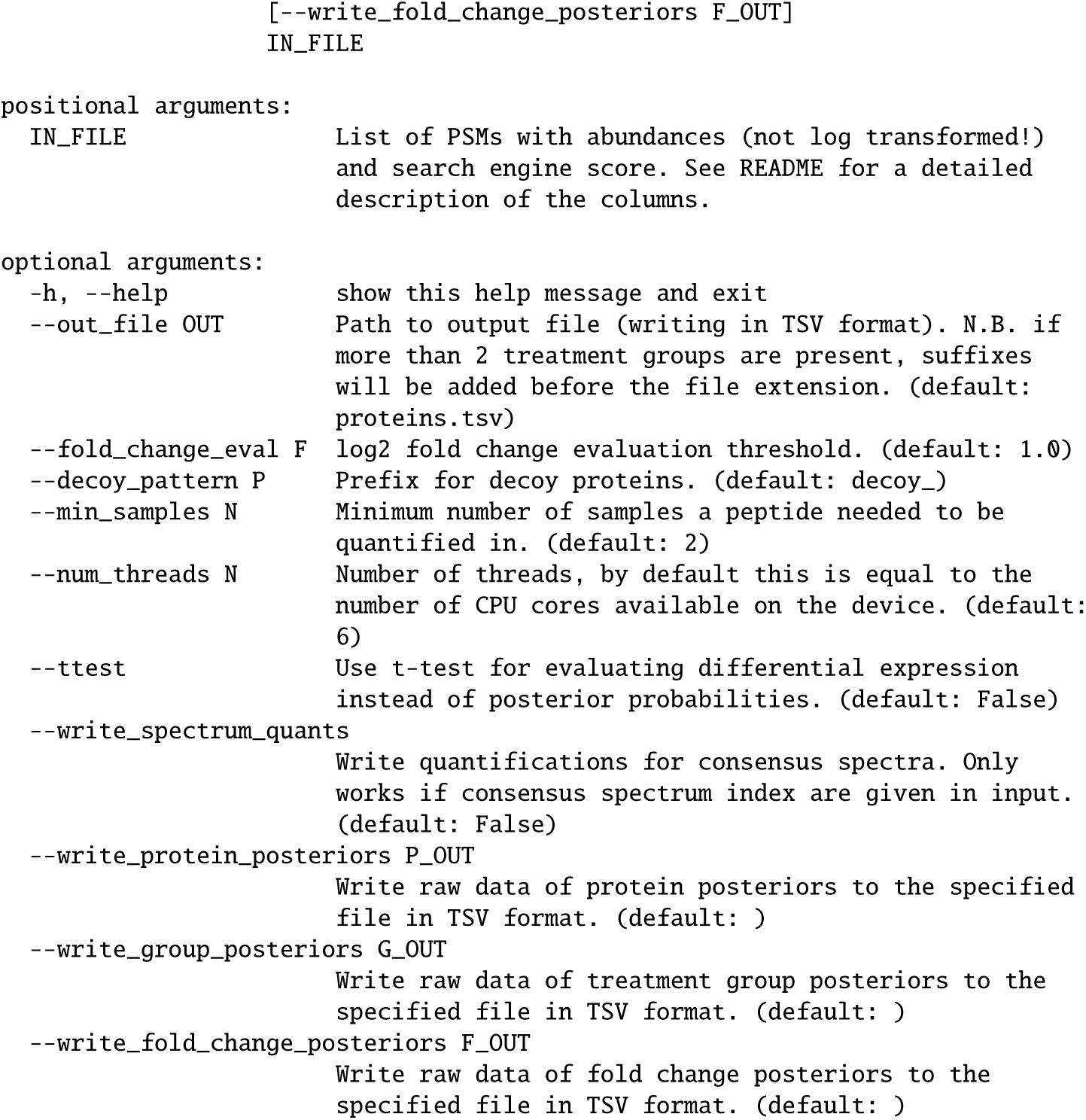

A detailed description of the command line arguments follows below:

~~~
    <IN_FILE>
~~~

tab-separated input file with the format described previously (*see* Subheading 2.6).

~~~
    --out_file <OUT>
~~~

tab-separated results file on protein-level. If more than 2 treatment groups are specified, multiple output files are generated, where the comparison is inserted before the `.tsv extension`. For example, if the output file is specified as `proteins.tsv`, then the output files will be named `proteins.1vs2.tsv, proteins.1vs3.tsv`, etc.

~~~
    --fold-change-eval <F>
~~~

the log_2_ fold change used for evaluation of differential abundance. Specifically, we integrate the probability of the fold change distribution outside the region [−*F, F*] as the probability that the protein is differentially abundant (*see* Subheading 3.7, **Step 9**). Note that this test is quite different from the *t*-test, and setting *F* = 0 to supposedly obtain all differentially abundant proteins regardless of the fold change will result in nonsensical results (*see* Subheading 3.7, **Step 10**).

~~~
    --decoy_pattern <P>
~~~

prefix for decoy proteins used by the search engine. For MaxQuant searches, this is typically `REV__.` This pattern is used to recognize which of the PSMs are targets and decoys.

~~~
    --min_samples <N>
~~~

this flag controls which peptides are discarded because they have too many missing values. For example, if *N* = 3 and a total of 10 samples are provided, all peptides with more than 7 missing values are discarded (*see* **Note 14**).

~~~
    --num_threads <N>
~~~

this flag controls how many threads are used to calculate the posterior distributions (*see* **Note 15**).

~~~
    --ttest
~~~

specifying this flag computes a *t*-test based on the expected values of the protein abundances instead of using the Bayesian posterior calculation. This is only meant for comparison purposes and we do not claim or support any validity of these results.

~~~
    --write_spectrum_quants
~~~

specifying this flag produces an extra intermediate output very similar to the peptides output specifically for Quandenser input. Instead of outputting the peptide abundances across all runs, it outputs the abundances for an MS1 feature group across all runs.

~~~
    --write_protein_posteriors <P_OUT>
~~~

writes the raw results of the posterior distributions of the relative protein abundance for each protein. How to visualize these results is described later (*see* Subheading 3.7, **Step 7)**.

~~~
    --write_group_posteriors <G_OUT>
~~~

writes the raw results of the posterior distributions of the treatment group mean abundance for each protein. How to visualize these results is described later (*see* Subheading 3.7, **Step 8)**.

~~~
    --write_fold_change_posteriors <F_OUT>
~~~

writes the raw results of the posterior distributions of the fold change for each protein. How to visualize these results is described later (*see* Subheading 3.7, **Step 9**).

### 3.5 Running Triqler

1. Download the example file from our GitHub repository: https://github.com/statisticalbiotechnology/triqler/blob/master/example/iPRG2016.tsv (*see* **Note 16**).
2. Run Triqler, keeping all parameters at their default values:

~~~
  $ python -m triqler iPRG2016.tsv
~~~

Below, the internal steps of Triqler and the produced outputs are explained section by section.
3. Boilerplate welcome message, including the specified command line parameters for easy future reference:

~~~
  Triqler version 0.5.0
  Copyright (c) 2018-2020 Matthew The. All rights reserved.
  Written by Matthew The (matthew.the@scilifelab.se) in the
  School of Engineering Sciences in Chemistry, Biotechnology and Health at the
  Royal Institute of Technology in Stockholm.
  Issued command: triqler.py iPRG2016.tsv
~~~
4. Triqler parses the input file and recalculates q-values and posterior error probabilities (PEPs) using a Python re-implementation of qvality [12].

~~~
  Parsing triqler input file
    Reading row 0
  Calculating identification PEPs
    Identified 12113 PSMs at 1% FDR
~~~
5. Some peptides may have been assigned to multiple MS1 features in a single run, each with a different XIC intensity (*see* **Note 17**). In this step, Triqler selects the best MS1 feature based on the search engine score and, if available, the feature-match error probability.

~~~
  Selecting best feature per run and spectrum
    featureGroupIdx: 0
~~~
6. Triqler divides the XIC intensities by a power of 10 while preserving 2 significant digits for the smallest observed intensity. This does not affect subsequent analysis nor the content of the output files but serves to increase the readability of the intermediate files (*see* **Note 18**).

~~~
  Dividing intensities by 100000 for increased readability
~~~
7. Peptide-intensity pairs from the different runs are grouped based on sequence and charge state. Subsequently, the FDR and PEPs of the unique peptides are calculated [13]. The grouped peptide-intensity pairs are then written to an intermediate file, in this case `iPRG2016.tsv.pqr.tsv` (*see* **Note 19**).

~~~
  Calculating peptide-level identification PEPs
    Identified 1988 peptides at 1% FDR
  Writing peptide quant rows to file: iPRG2016.tsv.pqr.tsv
~~~
8. Protein inference is executed using the best peptide score as the protein’s score, together with the picked protein approach [14].

~~~
  Calculating protein-level identification PEPs
    Identified 349 proteins at 1% FDR
~~~
9. Triqler makes use of the empirical Bayes method to estimate hyperparameters from the input data. We fit distributions to naive estimates of peptide and protein abundance values (*see* Subheading 3.8).

~~~
  Fitting hyperparameters
    params[“muDetect”], params[“sigmaDetect”] = 1.056334, 0.372395
    params[“muXIC”], params[“sigmaXIC”] = 3.276315, 0.953023
    params[“muProtein”], params[“sigmaProtein”] = 0.066437, 0.239524
    params[“muFeatureDiff”], params[“sigmaFeatureDiff”] = -0.013907, 0.149265
    params[“shapeInGroupStdevs”], params[“scaleInGroupStdevs”] = 1.027176,
  0.089433
~~~
10. The hyperparameters are now used in the probabilistic graphical model to estimate posterior distributions for the relative protein abundances, as well as for the fold change between the treatment groups.

~~~
  Calculating protein posteriors
    50 / 422 11.85%
    100 / 422 23.70%
    150 / 422 35.55%
    200 / 422 47.39%
    250 / 422 59.24%
    300 / 422 71.09%
    350 / 422 82.94%
    400 / 422 94.79%
~~~
11. Finally, the protein identification PEP and the probability that the fold change exceeds the fold change threshold specified by the --fold_change_eval parameter are combined. This combined PEP can then be used to compute the differential abundance FDR (*see* **Note 20**). For each comparison, Triqler then reports the number of differentially abundant proteins at 5% FDR and outputs the final results to one file per comparison (e.g. `proteins.1vs2.tsv`).

~~~
  Comparing 1:A+B to 2:B
    output file: proteins.1vs2.tsv
    Found 204 target proteins as differentially abundant at 5% FDR
  Comparing 1:A+B to 3:A
    output file: proteins.1vs3.tsv
    Found 216 target proteins as differentially abundant at 5% FDR
  Comparing 2:B to 3:A
    output file: proteins.2vs3.tsv
    Found 352 target proteins as differentially abundant at 5% FDR
  Triqler execution took 22.869995618006214 seconds wall clock time
~~~

In this particular dataset, there are a total of 383 truly differentially abundant (spiked-in) proteins. However, in the *A* + *B* vs *A* and *A* + *B* vs *B* comparisons, the expected log_2_ fold change coincides with the chosen `--fold_change_eval` parameter of 1.0, which makes it hard for about half of these proteins to be called differentially abundant (*see* **Note 21**).

### 3.6 Interpreting the Triqler output

The protein output files can be opened in Excel, or another spreadsheet package of choice. For an explanation of the columns *see* Table 1.

**Table 1.**
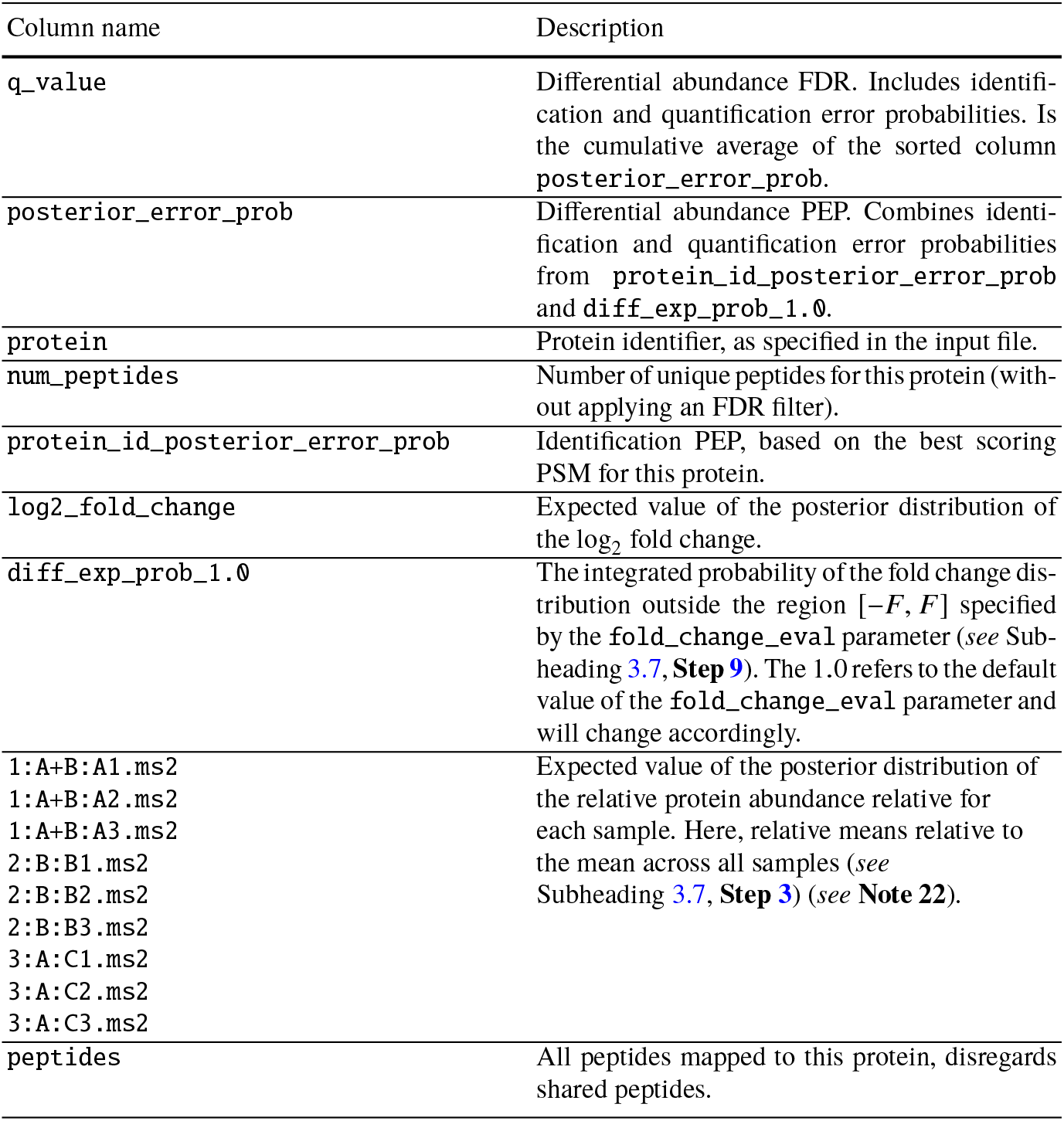
Triqler protein output format.

| Column name | Description |
| --- | --- |
| q_value | Differential abundance FDR. Includes identification and quantification error probabilities. Is the cumulative average of the sorted column `posterior_error_prob`. |
| posterior_error_prob | Differential abundance PEP. Combines identification and quantification error probabilities from `protein_id_posterior_error_prob` and `diff_exp_prob_1.0`. |
| protein | Protein identifier, as specified in the input file. |
| num_peptides | Number of unique peptides for this protein (without applying an FDR filter). |
| protein_id_posterior_error_prob | Identification PEP, based on the best scoring PSM for this protein. |
| log2_fold_change | Expected value of the posterior distribution of the $\log_2$ fold change. |
| diff_exp_prob_1.0 | The integrated probability of the fold change distribution outside the region $[-F, F]$ specified by the `fold_change_eval` parameter (*see* Subheading 3.7, **Step 9**). The 1.0 refers to the default value of the `fold_change_eval` parameter and will change accordingly. |
| 1:A+B:A1.ms2<br>1:A+B:A2.ms2<br>1:A+B:A3.ms2<br>2:B:B1.ms2<br>2:B:B2.ms2<br>2:B:B3.ms2<br>3:A:C1.ms2<br>3:A:C2.ms2<br>3:A:C3.ms2 | Expected value of the posterior distribution of the relative protein abundance relative for each sample. Here, relative means relative to the mean across all samples (*see* Subheading 3.7, **Step 3**) (*see* **Note 22**). |
| peptides | All peptides mapped to this protein, disregards shared peptides. |

To help understand the output better, we will look at a specific example in the `proteins.1vs2.tsv file`.

1. Search for the protein `HPRR3730445_poolB`, which should be around the 174th line.
2. In the 1st column, the *q*-value for this protein is shown to be 0.006985 (the exact value may vary in newer versions), meaning that the list of proteins up to this position has an FDR of 0.7%, meaning that we expect approximately 173 × 0.006985 = 1.2 false positives in this list of 173 proteins. This *q*-value combines the error probabilities of the identification and the quantification process, which improves on the typical practice of separately reporting identification and quantification FDRs.
3. In the 2nd column, we can see that the posterior error probability is 0.1302. This value, again, combines both identification and quantification error probabilities. Practically speaking this value means that this particular protein has a probability of 87% to be both correctly identified and differentially abundant.
4. In the 4th column, we see that we have found 3 unique peptides for this protein. However, this does not tell us how many were identified at 1% peptide-level FDR.
5. In the 5th column, the identification posterior error probability is listed as 0.0001162. Meaning that the protein has a probability of only 0.01% to be incorrectly identified.
6. In the 6th column, we see that the expected log_2_ fold change between the groups *A* + *B* and *B* for this protein is −1.522. This means that we expect that this protein is present in group *A* + *B* at a concentration of 2^−1.522^ = 0.34 relative to group B. According to the spike-in ratios, this value should have been 0.5.
7. In the 7th column, the probability that the log_2_ fold change between the *A* + *B* and *B* treatment group is larger than 1.0 (the default value for the `fold_change_eval` parameter) is given as 0.1301, meaning that there is a 13% chance that the absolute fold change was in fact below 0.5. If we would have specified a lower value for the `fold_change_eval` parameter, this probability would have increased, as we would be integrating a larger part of the posterior distribution of the fold change. This in turn will affect the combined posterior error probability and FDR in the first and second columns.
8. In the 8 – 16th column, we can see the individual expected protein abundance value per sample, as obtained from the posterior distributions for the relative protein abundance (relative to the mean over all samples, which would itself be represented by the value 1.0, also *see* Subheading 3.7, **Step 3**). For example, we see that in `A3.ms2` the expected value for the relative protein abundance is 5.475, meaning that it is 5.5 times higher than the average protein abundance across all samples. In `B2.ms2` this value is 13.64, meaning that we expect the abundance in this sample to be 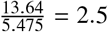 times higher than in `A3.ms2`.
9. In the 17th column and beyond, the 3 peptides that were used to identify and quantify this protein are listed.

### 3.7 Visualizing and interpreting posterior distributions

In some cases, one would like to have a closer look at how Triqler arrived at the conclusion of differential abundance for a particular protein. For this, Triqler provides functionality for extracting the relevant data for a protein of interest and plotting the posterior distributions at different levels (protein, treatment group, fold change).

1. Generate the posterior distribution plots for our protein of interest (we need to increase `--plot_max_fold_change` from its default value of 2.0, since we have extreme fold changes in this particular sample):

~~~
  $ python -m triqler.distribution.plot_posteriors --protein_id
  HPRR3730445_poolB --plot_max_fold_change 10.0 iPRG2016.tsv
~~~

Three plot windows will be opened, which we will examine shortly. First, we will look at the command line output.
2. After the boilerplate text and hyperparameter estimation, the peptides for this protein are listed in descending order of confidence. First, the “raw” intensity values are listed in the same order as in the regular protein output (see Table 1), note that these have been divided by the power of 10 as mentioned previously (*see* Subheading 3.5, **Step 6)** compared to the original Triqler input. Missing values are indicated by nan values. These are followed by the `combinedPEP`, which is a combination of the identification and feature-match error probability. In this case, since the latter is not included, `combinedPEP` just reflects the identification PEP.

~~~
  Peptide absolute abundances
  760.43 509.03 1028.25 842.80 1610.55 1289.44 nan nan nan
  combinedPEP=3.4e-06 peptide=R.WTAQGHANHGFVVEVAHLEEK.Q
  99.93 166.59 3184.98 1868.59 6260.46 5909.35 nan nan 59.71
  combinedPEP=2.2e-05 peptide=R.LVNQNASRWESFDVTPAVMR.W
  10064.52 12531.44 nan 27429.83 26226.20 23061.53 nan 242.17 19.53
  combinedPEP=0.0023 peptide=R.WESFDVTPAVMR.W
~~~

Here, we can see that all 3 peptides are identified with high confidence. There are many missing values in the group *A* samples (the last three columns of each row), and one missing value in the third *A* + *B* sample (third column).
3. The values from above are repeated, but now divided by the geometric mean across all samples, resulting in “relative” peptide abundances.

~~~
  Peptide relative abundances
  0.81 0.54 1.09 0.90 1.71 1.37 nan nan nan
  combinedPEP=3.4e-06 peptide=R.WTAQGHANHGFVVEVAHLEEK.Q
  0.12 0.21 3.96 2.32 7.78 7.34 nan nan 0.07
  combinedPEP=2.2e-05 peptide=R.LVNQNASRWESFDVTPAVMR.W
  2.70 3.37 nan 7.37 7.05 6.20 nan 0.07 0.01
  combinedPEP=0.0023 peptide=R.WESFDVTPAVMR.W
~~~
4. The above information (peptide intensities and identification PEPs) are then processed by Triqler’s probabilistic graphical model, resulting in several posterior distributions. First, the protein-level results are printed as expected values of the protein abundance posterior distribution. For comparison purposes, also the results of a regular *t*-test (2 treatment groups) or ANOVA test (3 or more treatment groups) on these expected values is shown.

~~~
  Protein abundance (expected value) and p-value
  3.27 2.97 5.48 7.66 13.64 11.55 0.01 0.07 0.02
  p-value: 3.282253815003453e-05
~~~
5. Next, Triqler calculates the probability that the log_2_ fold change is below the specified *--fold_change_eval* parameter. By default, this is 1.0, but we will explore how these probabilities change for different values, *see* Subheading 3.7, **Step 10.** We can see that for the *A* + *B* vs *B* comparison, there is a 13% probability that the log_2_ fold change is below 1.0, whereas for the other two comparisons, this probability is practically 0. This also makes sense, as in group *A*, this protein is in fact not present.

~~~
  Posterior probability |log2 fold change| < 1.00
    Group A+B vs Group B: 0.130072
    Group A+B vs Group A: 0.000000
    Group B vs Group A: 0.000000
~~~
6. Finally, we also summarize the posterior distributions of the group means by fitting normal distributions to them. These values are, again, relative to the mean across all group abundances. Note that these values are based on the log_10_ transformed values, rather than log_2_ as is the case for the values reported for fold changes. The standard deviations reflect the uncertainty in the estimates. In this case, the standard deviation for group *A* is the largest, which reflects the fact that the missing values cause a higher degree of uncertainty.

~~~
  Normal distribution fits for posterior distributions of treatment group relative abundances:
    Group A+B: mu, sigma = 0.472918, 0.114328
    Group B: mu, sigma = 0.918317, 0.071100
    Group A: mu, sigma = -1.782820, 0.269253
~~~
7. Now, we can examine the 3 posterior distribution plots, starting with the relative protein abundance distributions in Fig. 3. Note that the abundance values are, again, log_10_ transformed. The distributions for the samples in the group *A* + *B* and *B* are relatively narrow, due to the high number of observed values, in contrast to the samples in group *A*, which have a large number of missing values combined with very low intensities.
8. For the posterior distributions of the group means, we can clearly see the effect of the individual samples from the previous step (Figure 4). Here, we can also verify that the distributions do resemble normal distributions, as was assumed earlier (*see* Subheading 3.7, **Step 6**).
9. Finally, for each pair of groups, we “subtract” one group mean posterior distribution from the other to obtain a fold change posterior distribution (Figure 5). Again, note the logarithm change from log_10_ to log_2_ compared to the previous step. In these violin plots, we show the distribution on the *y*-axes. The green distributions represent the Triqler estimations and, for comparison, we also display the estimate that would have been given by a naive method (Top 3, mean row imputation) in blue.
10. To show the effect of the `--fold_change_eval` parameter, we now change this to 0.5 instead of the default value of 1.0.

~~~
  $ python -m triqler.distribution.plot_posteriors --protein_id
  HPRR3730445_poolB --plot_max_fold_change 10.0
  --fold_change_eval 0.5 iPRG2016.tsv
~~~ All of the command line output stays the same, except for the second to last block, where the *A* + *B* vs *B* comparison now also shows a very low probability of being below the lowered `--fold_change_eval.`:

~~~
  Posterior probability |log2 fold change| < 0.50
    Group A+B vs Group B: 0.010441
    Group A+B vs Group A: 0.000000
    Group B vs Group A: 0.000000
~~~

**Fig. 3.**
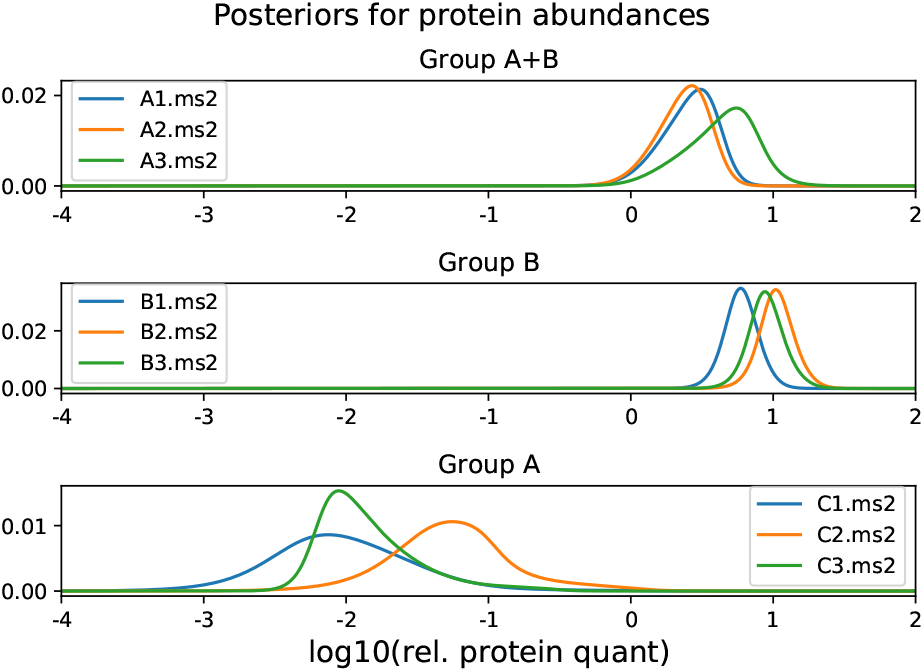
Posterior distributions for the relative protein abundance of HPRR3730445_poolB in each of the 9 samples.

**Fig. 4.**
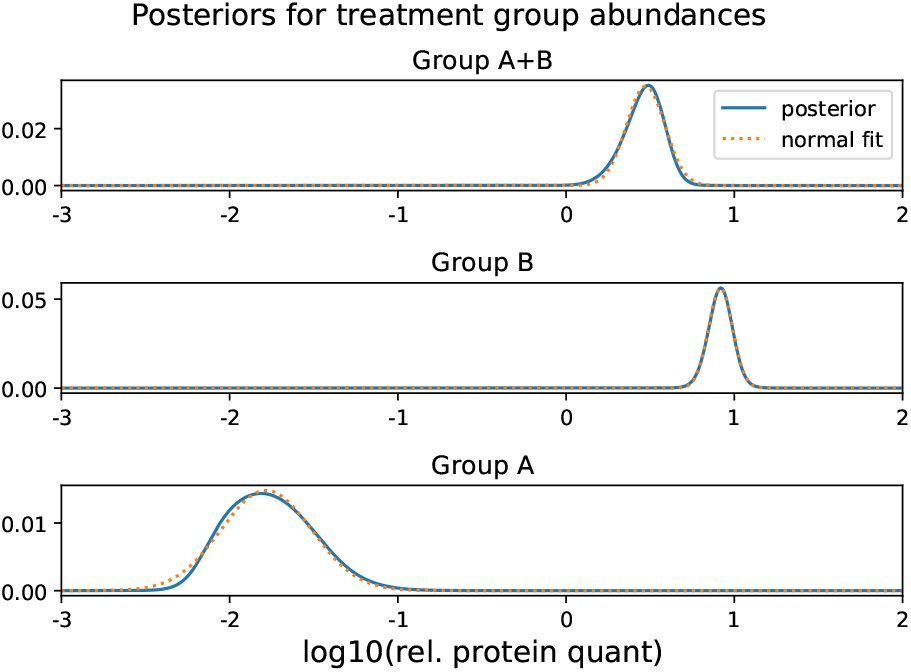
Posterior distributions for the relative mean group abundance of HPRR3730445_poolB in each of the 3 groups.

**Fig. 5.**
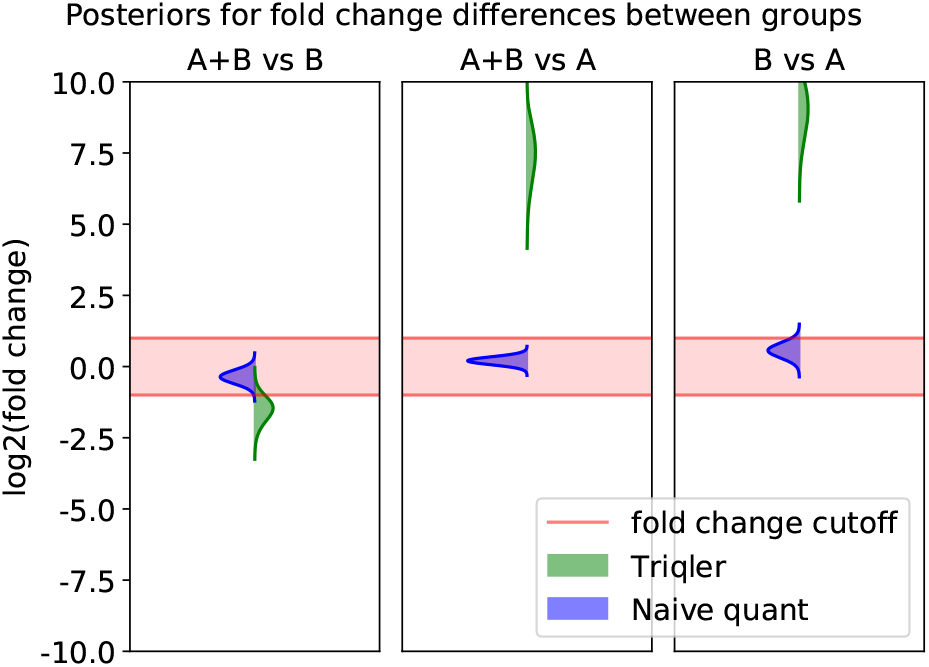
Posterior distributions for the log2 fold changes of HPRR3730445_poolB for each of the 3 group comparisons.

From the new fold change posterior distribution plot (Figure 6), we can also see that the distribution for the *A* + *B* vs *B* comparison now is practically completely outside of the red fold change region. At the same time, this demonstrates the danger of using a too low value for `--fold_change_eval`, as the full width of the green distribution is now larger than the red region. Even if the green distribution would be centered around 0, there would still be some probability outside of the red region, which would then be reported as a non-negligible probability of the protein being differentially abundant even though no actual difference can be seen.

**Fig. 6.**
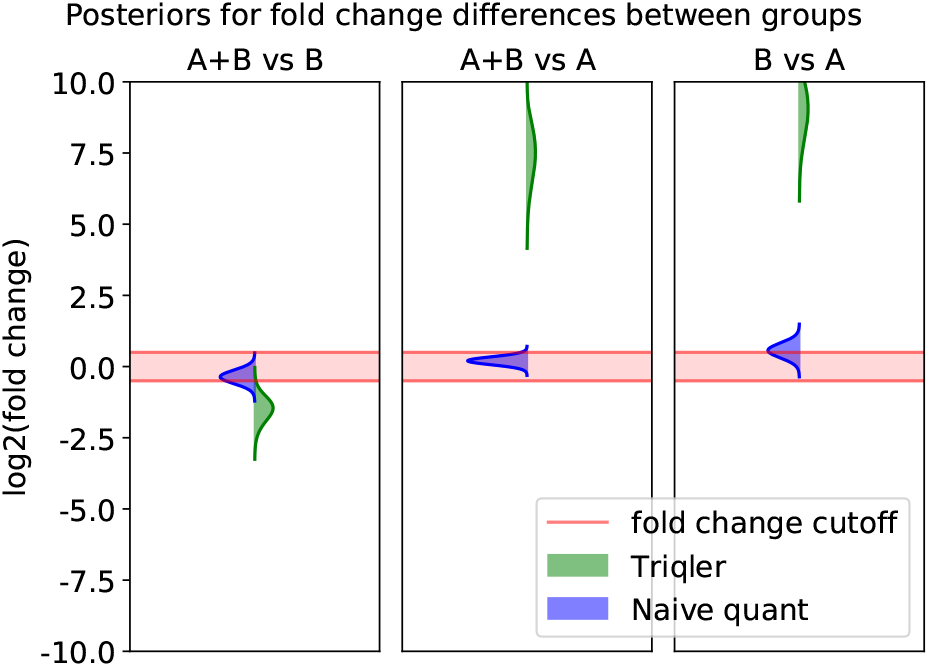
Same as Fig. 5 but with the log_2_ fold change evaluation threshold changed to 0.5 instead of the default value of 1.0.

### 3.8 Visualizing and interpreting hyperparameter estimation

A common concern for using Bayesian methods is the dependence on the prior distributions. Triqler employs the Empirical Bayes method to estimate the hyperparameters for the prior distributions (*see* **Note 23**) for the probabilistic graphical model. To check if these hyperparameter estimations are reasonable, Triqler provides functions to investigate these through graphical inspection (*see* **Note 24**).

1. Generate the fits for the hyperparameter estimations:

~~~
  $ python -m triqler.distribution.plot_hyperparameter_fits iPRG2016.tsv
~~~
2. To estimate the probability of a missing value as a function of the XIC, Triqler postulates that the log_10_(XIC) of all peptides across all samples can be modeled as a left-censored normal distribution (Fig. 7). Here, the left-censored normal distribution (green) is a normal distribution (cyan) with some “mass” missing for low XICs due to a sigmoidal censoring function (purple). Note that the XIC values have been divided by a power of 10 (*see* Subheading 3.5, **Step 6)**. Note that, in this particular example, the influence of the sigmoidal censoring function is not very apparent, likely due to the design of the study with many actually missing values.
3. To estimate the prior distribution for the relative protein abundances we fit a hyperbolic secant function (*see* **Note 25**) to naive estimations (simple average of relative peptide abundances) of the protein abundances (Fig. 8).
4. To estimate the distribution of the difference between the true and observed XIC, we fit a hyperbolic tangent distribution to the difference between the observed XICs and the expected XIC based on the relative protein abundance estimated in the previous step and the estimated ionization efficiency (*see* **Note 26**). (Fig. 9). This can be seen as an estimate for the measurement uncertainty which will then propagate through the graphical model.
5. Finally, we estimate the distribution of the standard deviation between protein abundances of samples within the same treatment group using a Gamma distribution (Fig. 10). This serves to capture the biological and/or technical variance within a group.

**Fig. 7.**
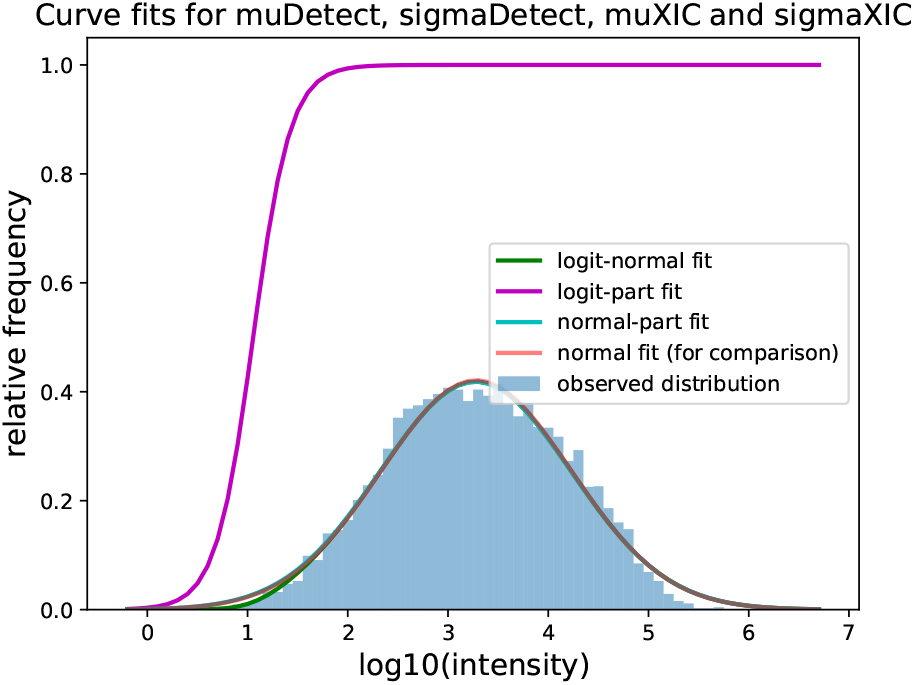
Estimating the hyperparameters for the missing value probability as a function of XIC by a left-censored normal distribution.

**Fig. 8.**
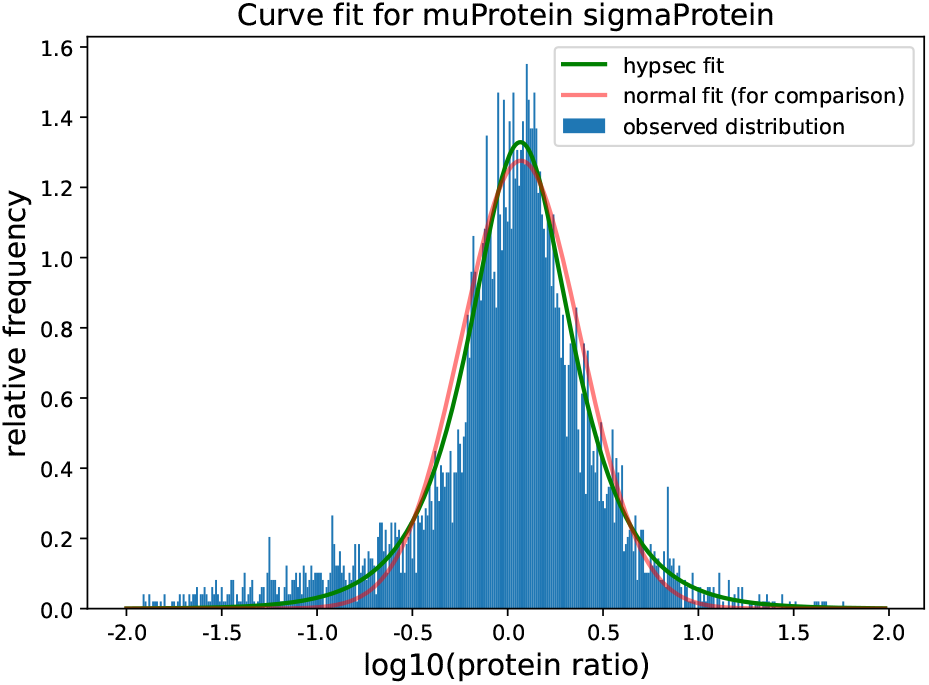
The prior for the mean of the relative protein abundance is estimated with a hyperbolic secant distribution.

**Fig. 9.**
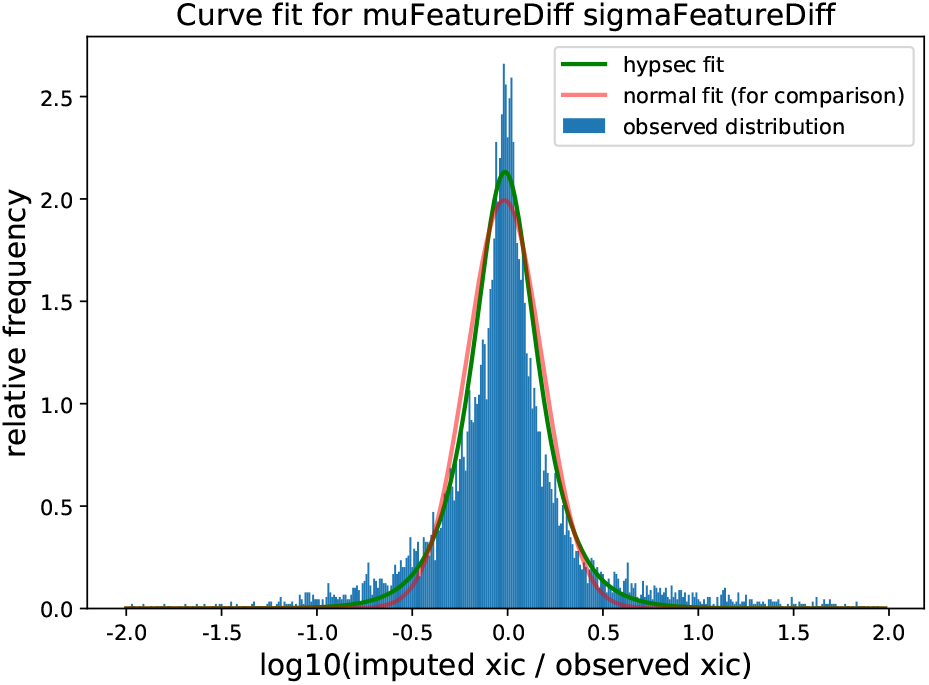
The measurement uncertainty distribution is estimated with a hyperbolic secant distribution.

**Fig. 10.**
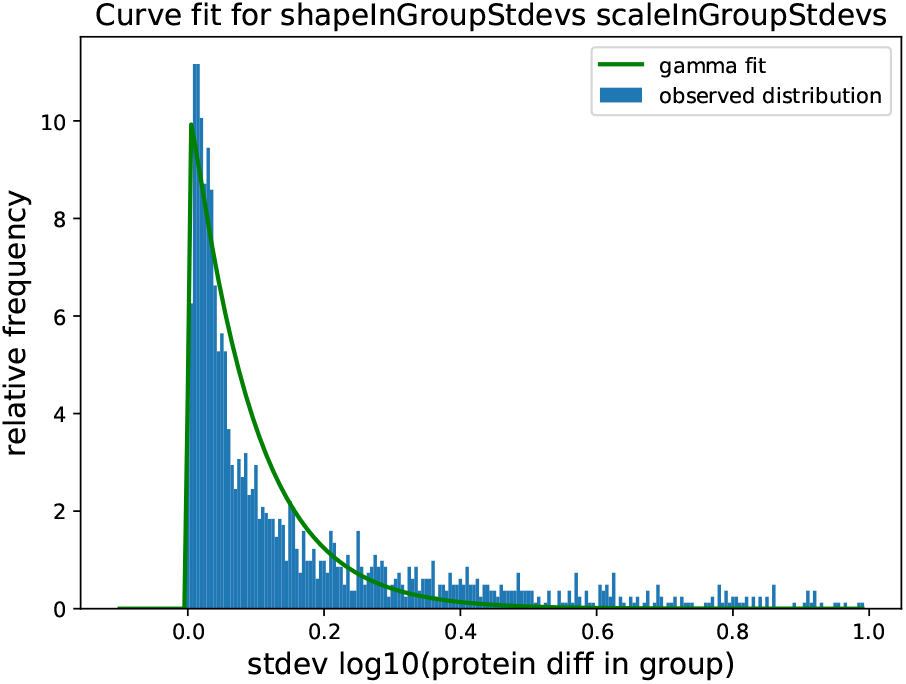
The within-group standard deviation distribution is estimated by a Gamma distribution.

## Notes

1. We have, however, observed that Triqler still is able to correctly distinguish between changing and unchanging proteins in engineered datasets if the former requirement is violated.
2. Currently, shared peptides are not considered for identification and quantification. However, we expect to include functionality to include shared peptides in the near future.
3. For some test datasets, the results look reasonable for the default thresholds of 1% FDR as well, but as the error estimates are likely unreliable we do not recommend running Triqler in this way.
4. We have verified the validity of the results without MBR on several test datasets. Triqler will work with match-between-runs turned on, but we cannot guarantee the validity of the results. In a future version, we will try to incorporate the error estimates from MaxQuant’s MBR step, similar to what we have done with Quandenser (*see* Subheading 3.2).
5. If the output from `crux percolator` is used, make sure that the order of the files processed by crux `tide-search` is the same as in the file specified at `--file_list_file`.
6. A more detailed example of how to run Quandenser and obtain search results from the consensus spectra can be found here: https://github.com/statisticalbiotechnology/quandenser/wiki/Example:-Quandenser-followed-by-Tide-and-Triqler
7. Currently, this list includes SEQUEST [15] (Comet [16], Tide [17]), MSGF+ [18], X!Tandem [19]. Custom Python scripts are available for MODa [20], MSFragger [21] and Andromeda [22] upon request to the authors.
8. *see* **Note 7.**
9. This can be installed through `$ pip install simsalabim`. Also, *see* **Note 10.**
10. A more detailed example of how to run the dinosaur adapter and obtain search results can be found here: https://github.com/MatthewThe/simsalabim/wiki/Example:-Dinosaur-followed-by-Tide-and-Triqler
11. *see* **Note 10.**
12. We strongly recommend searching the recalibrated MS2 spectrum files, which now have accurate MS1 precursors assigned. This generally improves the identification rate and allows for multiple identifications per spectrum for chimeric spectra. In the dataset we used as an example in this manuscript (iPRG2016), we could increase the number of identified peptides by 34%.
13. *see* **Note 7.**
14. We have observed in [7] that even for low values of *N*, e.g. *N* = 3, the differential abundance FDR remains under some form of control, but advise users to be careful with setting this value so low, as the number of false positives does increase.
15. On some systems, the python multiprocessing module causes issues. Setting *N* = 1 will bypass this multiprocessing module at the cost of longer runtimes.
16. This dataset is described in [23]. Briefly, this dataset was designed to test how well protein identification and quantification pipelines can deal with shared peptides, by using two pools of synthesized proteins which are similar to human proteins (PrESTs). Each of the pools contained one out of a pair of proteins that share a number of peptides. In the first 3 samples, both pools were mixed in equal parts into a background of E. Coli lysate, in the second 3 samples, only the B pool proteins were mixed in, and in the last 3 samples, only the A pool proteins were added.
17. This can, for example, be a result of problems with MS1 feature detection or due to false positives in the MS2 spectrum identification
18. Specifically `example/iPRG2016.tsv.pqr.tsv`, the output of the `--write-spectrum-quants` file, `example/iPRG2016.tsv.sqr.tsv` and the command line output of the `plot_posteriors` commands.
19. This file summarizes the Triqler input file by grouping the rows by combinations of peptide and charge. The format consists of a column called combinedPEP, which combines the peptide-level identification PEP and the feature-match PEP. In the absence of the latter, it is just the identification PEP. After the charge column and two columns specific to Quandenser (feature group index and consensus spectrum scan number), there are three groups of *N* columns, where *N* is the number of samples. The first group contains the feature-match PEPs, the second group the intensity values (divided by the power of 10 as mentioned earlier) and the last group contains the identification PEP. The last two columns contain the peptide sequence and protein identifier respectively.
20. This is done by sorting the PEPs in ascending order and taking the cumulative average [24]. This works because the PEP is the derivative of the FDR, and is therefore sometimes called the local-FDR.
21. The 192 poolA proteins and 191 poolB proteins are present at half the concentration in the *A* + *B* samples compared to the *A* and *B* samples respectively. The posterior distribution therefore, ideally, centers around a log_2_ fold change of 1.0. Here, half of the probability distribution would indicate a log_2_ fold change below 1.0 resulting in a posterior error probability of 0.5. In practice, the log_2_ fold change posterior distributions should include the true log_2_ fold change of 1.0, but do not necessarily center around 1.0. Therefore, some of these proteins will have a fold change posterior distributions centered above 1.0 and a posterior error probability < 0.5, which might result into them being called significantly differentially abundant at 5% FDR.
22. Another way to consider these values is as the regular protein abundance values (e.g. summed intensity) divided by the mean over all samples.
23. For a short introduction for non-statisticians into the basics of Bayesian statistics, including priors, hyperparameters, posteriors and some simple examples, see [25].
24. See the Supplemental Data of [5] for more examples of fitted hyperparameter distributions for a range of datasets
25. The hyperbolic secant distribution (green) has heavier tails than the normal distribution (red) and is, thus, better able to deal with outliers that are common in real-world data.
26. The ionization efficiency is estimated as the average across all samples of the XIC divided by the relative peptide abundance, where missing values are not considered.

